# Heritable variation in resistance to the endonuclear parasite *Holospora undulata* across clades of *Paramecium caudatum*

**DOI:** 10.1101/2020.09.08.288118

**Authors:** Jared Weiler, Giacomo Zilio, Nathalie Zeballos, Louise Noergaard, Winiffer D. Conce Alberto, Sascha Krenek, Oliver Kaltz, Lydia Bright

## Abstract

Resistance is a key determinant in interactions between hosts and their parasites. Understanding the amount and distribution of genetic variation in this trait can provide insights into (co)evolutionary processes and their potential to shape patterns of diversity in natural populations. Using controlled inoculation in experimental mass cultures, we investigated the quantitative genetic variation in resistance to the bacterial parasite *Holospora undulata* across a worldwide collection of strains of its ciliate host *Paramecium caudatum*. We combined the observed variation with available information on the phylogeny and biogeography of the strains. We found substantial variation in resistance among strains (with broad-sense heritability > 0.5), repeatable between laboratories and ranging from total resistance to near-complete susceptibility. Early (one week post inoculation) measurements provided higher estimates of resistance heritability than did later measurements (2-3 weeks), possibly due to diverging epidemiological dynamics in replicate cultures of the same strains. Genetic distance (based on a neutral marker) was positively correlated with the difference in resistance phenotype between strains (r = 0.45), essentially reflecting differences between highly divergent clades (haplogroups) within the host species. Haplogroup A strains, mostly European, were less resistant to the parasite (49% infection prevalence) than non-European haplogroup B strains (28%). At a smaller geographical scale (within Europe), strains that are geographically closer to the parasite origin (Southern Germany) were more susceptible to infection than those from further away. These patterns are consistent with a picture of local parasite adaptation. Our study demonstrates ample natural genetic variation in resistance on which selection can act and hints at symbiont adaptation producing signatures in geographic and lineage-specific patterns of resistance in this model system.

## 1 Introduction

Evolutionary interactions between hosts and their symbionts can shape patterns of genetic diversity and adaptation in natural communities (Kaltz 1998; Howells *et al*. 2013; Caldera *et al*. 2019). For this diversification to occur there must be heritable variation in the traits determining the ‘compatibility’ between a host and a symbiont, from resistance and infectivity to more complex life-history or developmental traits (Law 1998; Dale and Moran 2006; Chomicki 2020). In many systems, this variation can be assessed in controlled infection experiments, using different combinations of host and symbiont genotypes (Carius 2001; Kaltz 2002; Cayetano and Vorburger 2013). Such experiments not only inform on the present evolutionary potential of the interacting players, but can also reveal patterns shaped by their (co)evolutionary past, such as local adaptation (Greischar and Koskella 2007; Hoeksema and Forde 2008), lineage or species specificity (Summerer *et al*. 2007; Chappell and Rausher 2016) and symbiont life styles (Degnan and Moran 2008; Sachs *et al*. 2011).

The ciliated protist *Paramecium* spp. is a useful potential model organism for experimental investigations into diverse associations with endosymbiotic bacteria that represent the full continuum of endosymbiotic interactions: from facultative to obligate, and from horizontal to mixed to vertical transmission (Preer *et al*. 1974; Fokin 2009; Fujishima 2009; Görtz 2009; Goertz 2010; Fujishima and Kodama 2012; Plotnikov *et al*. 2019). The possession of these endosymbiotic partners is likely the result of *Paramecium*’s bacterivorous nature, whereby some of its prey have evolved means to evade digestion and to take up residence in the cytoplasm of the cell, and also in the host nuclei (Goertz 2010; Boscaro *et al*. 2013; Schulz and Horn 2015. For many of these associations, the biology and evolution are still poorly understood. Among the notable exceptions is the genus Holospora, a group of Alphaproteobacteria distantly related to other endosymbionts, such as Rickettsia or Wolbachia (Szokoli *et al*. 2016; Garushyants *et al*. 2018; Potekhin *et al*. 2018; Munoz-Gomez *et al*. 2019). Following the discovery of *Holospora* bacteria in the 19th century (Hafkine 1890), there has been extensive research on their morphology, infection life cycle, and taxonomic relationships (for a historical overview, see (Fokin 2009)). More recently, the *Paramecium-Holospora* system has served as a model to study epidemiology and evolution in experimental microcosms (Lohse *et al*. 2006; Nidelet *et al*. 2009; Duncan *et al*. 2011a; Castelli 2015; Duncan *et al*. 2015; Dusi *et al*. 2015; Duncan 2018; Nørgaard 2020; Zilio 2020b; Zilio 2020a).

*Holospora* are highly species-specific endonuclear symbionts of *Paramecium* spp, with each species specifically infecting either the somatic macronucleus (MAC) or the germline micronucleus (MIC) of the host cell (Fokin 2009; Fujishima 2009). *Holospora* species infect one single host species, and only the phylogenetically more distant *Holospora caryophila* (now *Ca. Preeria caryophila*) is known to occur on multiple host species (Potekhin *et al*. 2018). Furthermore, the interaction is obligate for the bacterium, and replication is only possible inside the host cell, as many biosynthetic metabolic pathways are fractured or even missing (Fokin 2009; Garushyants *et al*. 2018). Reproductive forms of the bacteria (RFs) can be vertically transmitted from mother to daughter nuclei during host cell division (Fujishima 2009; Magalon *et al*. 2010), whereas horizontal transmission occurs when infectious forms (IFs) are released during cell division or upon host death and then ingested by new hosts while feeding. *Holospora* symbionts show typical parasite features: they consume resources (e.g., nuclear proteins and nucleotides) in vital host compartments, disrupt sexual processes (Görtz and Fujishima 1983) and reduce asexual division, cell survival, and motility (Restif 2006; Duncan *et al*. 2010; Bella 2016; Nørgaard 2020; Zilio 2020b). Several host resistance mechanisms have been described (Rautian 1993; Skoblo 1996; Fokin 1997; Fokin *et al*. 2003; Goertz 2010); they include reduced uptake of IFs (Fels *et al*. 2008), inability (of the bacteria) to enter the nucleus (Rautian 1993) or being expelled from the nucleus after entry and before differentiating into RFs (Rautian 1993; Fokin 2005), and/or surveillance and lysis of the bacteria after it has entered and taken up residence in the nucleus (Fokin 1997; Skovorodkin 2001; Goertz 2010).

Little is known about *Paramecium-Holospora* interactions in natural populations. *Holospora* have been found at low infection prevalences in temperate or elevated locations around the globe (Fokin 1993; Hori and Fujishima 2003; Fokin *et al*. 2004; Fokin 2006; Serra *et al*. 2016), but can occasionally cause local epidemics (Fokin 2009), which might give a strong selective advantage to resistant variants over susceptible ones. Previous cross-inoculation studies demonstrated the existence of such natural variation, showing that different strains of *Paramecium caudatum* (and several other species) have different qualitative infection phenotypes, with some strains appearing universally susceptible to infection with *Holospora*, while others are more difficult to infect or even entirely resistant (Fujishima and Fujita 1985; Rautian 1993; Rautian 1996; Skoblo 1996; Potekhin *et al*. 2018). Previous studies have also compared resistance across independent mating groups or ‘syngens’. Syngens have long been known for *Paramecium* (Gilman 1941), and although reproductive isolation may not always be complete (Tsukii and Hiwatashi 1983; Johri *et al*. 2017), genetic analyses identified various clades in *P. caudatum* that can be considered as independent evolutionary units (IEUs) (Barth *et al*. 2006; Hori *et al*. 2006; Johri *et al*. 2017; De Souza 2020). These clades diverged up to 20 MYA ago (De Souza 2020) and potentially represent ‘cryptic species’ that are morphologically identical, but genetically isolated. Given the strong species-specificity of *Holospora*, such cryptic species complexes may represent an important structural component of resistance variation in natural populations. Indeed, Fujishima and Fujita (1985) found that *Holospora obtusa* was only able to infect strains from 5 out of 6 of the *P. caudatum* syngens studied, whereas an extensive study on *Holospora undulata* revealed successful infection of 84 of 92 *P. caudatum* strains and all 10 syngens tested (Skoblo 1996). In yet another system, Rautian et al (1993) reported evidence of strong syngen-level specificity of strains from *Holospora acuminata* on *Paramecium bursaria*, and this across a wide range of geographic origins.

However, while most of the classic studies have investigated the qualitative aspects of resistance, it clearly is a continuous trait: inoculation of experimental cultures typically results in higher or lower proportions of infected individuals (Lohse *et al*. 2006; Fels *et al*. 2008). Whether this reflects a polygenic basis of the trait or is simply due to the probabilistic nature of the infection process is unclear, but obviously continuous trait variation may produce very different epidemiological or evolutionary dynamics than qualitative variation (Duncan *et al*. 2011b).

Our goal in this study was to provide a rigorous analysis of the amount and distribution of quantitative genetic variation in resistance in the *Paramecium*-*Holospora* system. We performed a resistance assay testing 30 *P. caudatum* strains from a worldwide collection against the *H. undulata* reference strain (Dohra 2013). Using basic statistical tools, we estimated the heritable fraction of the observed variation in resistance and compared this variation between highly divergent clades, mainly from two haplogroups (Europe vs. rest of world). We further provide a first set of tentative analyses linking phylogenetic distances (based on a neutral marker), geographic distances and trait variation among the strains.

## 2 Materials and methods

### 2.1 *Paramecium-Holospora* life cycle

The infection life cycle (Figure 1) is well-characterized for different *Holospora* species (Görtz 1980; Fujishima 2009; Potekhin *et al*. 2018). All species show the same morphological and functional dimorphism, with infectious forms (IFs; elongated, up to 15 μm) and reproductive forms (RFs; more rounded, oblong, 5 μm). A horizontally acquired infection begins when the immobile IFs are taken up by a *Paramecium* in the course of feeding through the oral apparatus, and from there reach the nascent phagosome, or food vacuole (Figure 1A). For successful infection, the IFs must escape the quickly acidifying food vacuole, with the aid of an electron-translucent tip (Görtz *et al*. 1990; Dohra 1999; Fujishima 2009) and reach the cytoplasm by 30 minutes to one hour post-uptake.

**Figure 1.**
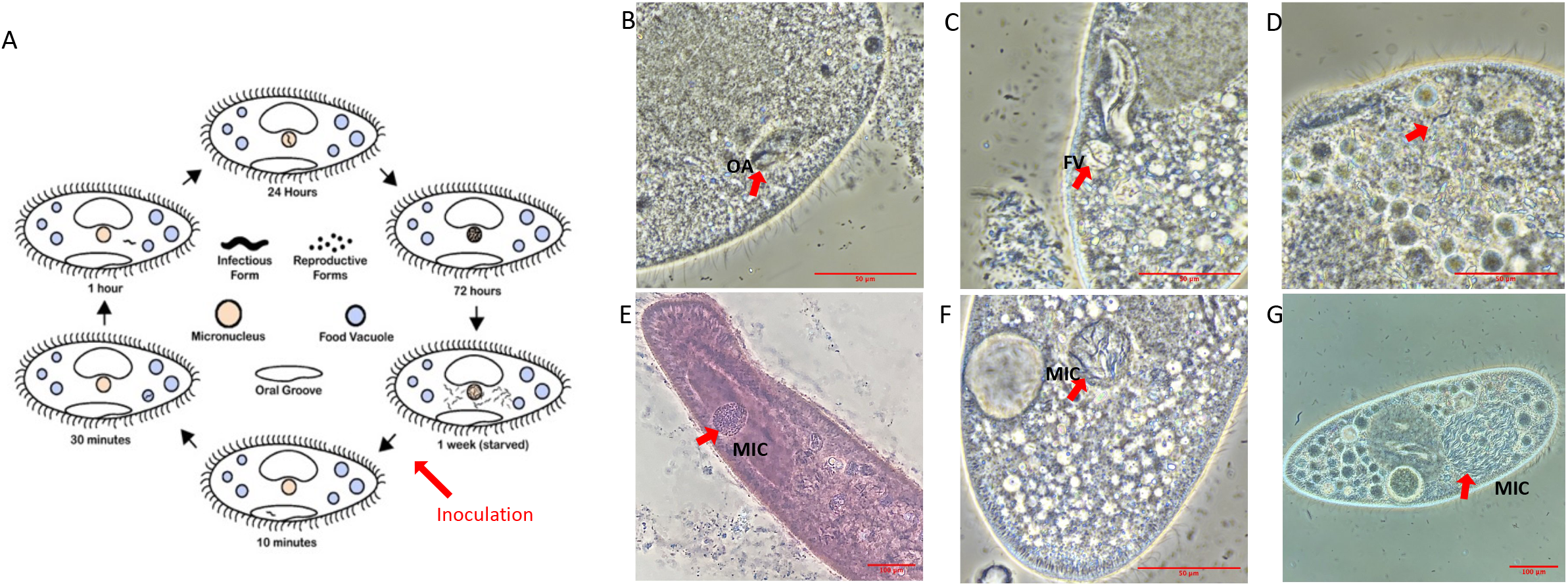
**A**. *Holospora* life cycle, diagrammed. **B-G**. Representative images of timepoints in infection of *Paramecium caudatum* by *Holospora undulata;* **B**. IF within the oral groove (OA) at 10 minutes p.i. **C**. IF in a food vacuole (FV) at 30 minutes p.i. **D**. IF in the cytoplasm at 1 hour p.i. **E**. RFs in the micronucleus (MIC) at 3 days p.i. (d.p.i.) **F, G**. IFs in the MIC at 7 d.p.i. **B-D, G**. Live cell images, differential interference contrast. **E**. Cell fixed with 5% glutaraldehyde and stained with lacto-aceto-orcein. **F**. Cells fixed with 5% glutaraldehyde. All cells imaged on a Nikon Ti-S inverted microscope, using phase contrast, 40X objective.

Between 1 hour and 24 hours post uptake, the *H. obtusa* IF has recruited host actin in order to drive its movement across the cell (Sabaneyeva *et al*. 2009) to the MAC and entered the nuclear envelope. This process involves an IF-specific protein, which has two predicted actin-binding domains and translocates to the outside of the bacterial cell during invasion (Iwatani *et al*. 2005). The same invasion tip seems to be used to exit the food vacuole (Figure 1B-D). The IFs then differentiate into the replicating RFs, thus forming characteristic clusters of bacterial cells in the nucleus. Replication continues over the first week post-infection, and the enlarging nucleus makes it easy to identify infected cells under the microscope at 400-1000x magnification (Figure 1E). After c. 1 week post infection, a developmental switch, possibly induced when a certain bacterial cell density is reached (Fujishima 1990; Kaltz and Koella 2003; Nidelet *et al*. 2009), causes RFs to differentiate into IFs, thereby further enlarging the nucleus (Figure 1F and G). Eventually, the IFs are released, either during host cell division or upon host death, thus completing the cycle.

### 2.2 *Paramecium caudatum* strains and culturing

The *P. caudatum* strains were taken from a collection with worldwide distribution (Table S1) at the Institute of Hydrobiology (TU Dresden, Germany). We deliberately chose strains across several clades (Tarcz *et al*. 2013; Johri *et al*. 2017), mostly from the A and B clades, and one each from the C and D clades, respectively. All strains had been uninfected at the time of collection.

A mix of infected long-term cultures were used to extract parasites for the resistance assays; these cultures represented several *P. caudatum* strains (of unknown COI genotype), all different from the 30 tested strains. Infections in all cultures originated from the same parasite strain, HU1 (Dohra 2013), isolated from a pond near Stuttgart (Germany) by H.-D. GÖrtz in 2000 and maintained in the OK lab since 2001.

All *P. caudatum* stocks were grown in organic lettuce medium inoculated with *Serratia marcescens* in the OK lab. Naïve and infected cultures were then transferred to Lydia Bright’s (LB; State University NY, US) lab, and kept in wheat grass medium inoculated with *Klebsiella* (Aury et al. 2006). For extended period storage, cultures were stored at 15-18° C; amplification of cultures and resistance assays were conducted at 23°C (OK lab) or at room temperature (22-25°C; LB lab).

### 2.3 Resistance assays

#### Inoculation

Resistance assays were conducted both in the OK lab (30 strains) and in the LB lab (18 strains), in the same year (2019; Table S1). To measure resistance, replicate cultures of a given strain were confronted with an inoculum of infectious forms, based on standard protocols, e.g. (Magalon *et al*. 2010). Briefly, large volumes of a highly infected *Paramecium* culture (up to 400 ml; preferably starved to induce the differentiation of infectious forms (Kaltz and Koella 2003), were concentrated by centrifugation, and all cells were then crushed mechanically in a bead beater to extract the infectious forms. The concentration of IFs was determined with a hemocytometer at 200X magnification under the microscope. In the OK lab, 10 mL samples from a given *P. caudatum* strain replicate (at carrying capacity) were gently centrifugated (700g for 15 min) and c. 1.5 mL of concentrated culture (∼3000-5000 host cells) were recovered. Then c. 10^4^ IFs from a freshly prepared inoculum were added to each replicate. Similarly, in the LB lab, 1-mL replicates of concentrated host cells (≤ 1.6 ⨯ 10^5^ cells) were inoculated with up to 10^5^ IFs. Final IF concentrations varied between experimental blocks (see below) but were the same for all replicates in a given block. On day 3 post inoculation (p.i.) in LB lab assays and day 4 p.i. in OK lab assays, we added 5 mL of medium to the inoculated replicates to prevent mortality due to high density. Additional medium (10-20 mL) was added after day 7 in order to maintain the replicates at carrying capacity.

#### Imaging and infection success

Infection success was measured by determining the number of infected paramecia from samples of 20-30 individuals from each replicate, after lacto-aceto-orcein fixation (GÖrtz 1980) and inspection of the individuals at 400-1000x magnification (phase contrast). The proportion of infected cells within a sample will be referred to as infection prevalence. Infection prevalence was measured at two time points. ‘Early’ measurements were taken on day 5 or 7 post inoculation. Most, if not all, infections establish during the first 48 h post inoculation (Fels et al 2008); over the following days infection prevalences remain stable and RFs multiply in the micronucleus, making infected cells easy to spot under the microscope (Fels and Kaltz 2006; Fels *et al*. 2008). Therefore our early measurements represent a good quantitative estimate of host resistance, here defined as the proportion of uninfected hosts (1 - infection prevalence) (Fels and Kaltz 2006; Nidelet *et al*. 2009; Magalon *et al*. 2010). ‘Late’ measurements of infection prevalence were taken on day 14 (LB lab) or 21 p.i. (OK lab), when infections have long taken over the micronucleus. At this point, the first infected cohort has already started to produce new secondary infections (typically during the second week p.i., (Duncan *et al*. 2015; Nørgaard 2020) and infected hosts also show reduced division rate and increased mortality, leading to more complex natural epidemiological dynamics in the culture (Restif 2006; Fokin 2009; Fujishima 2009; Magalon *et al*. 2010). Hence, late infection prevalences are likely determined by multiple traits, and not just host resistance.

#### Experimental replication

For the OK lab assays, two independent experimental blocks were established one week apart. For each block, three *Paramecium* replicate cultures per strain were grown to carrying capacity over one week prior to inoculation. One single inoculum was prepared for each block, which was then distributed over the three replicates, as described above (2 blocks x 30 strains x 3 = 180 inoculated replicates). Early measurements were taken for 177 replicates (30 strains) on day 5 p.i.; 116 late measurements (21 strains) were taken on day 21 (block 1) and day 14 (block 2). The LB lab assays (18 strains) were carried out in 4 independent blocks, over the course of 6 weeks, with a total of 38 inoculated replicates (1-4 replicates per strain; median = 2). As in the OK lab assays, independent replicate cultures and inocula were used for each block. Early measurements were taken on day 5 p.i. and day 7 p.i. (67 observations) and late measurements on day 14 p.i. (38 observations). See Table S1 for a summary of the replicates per strain and lab.

### 2.4 Genotyping and phylogeny

The relatedness of the stocks of *P. caudatum* was determined from single marker genotyping at the cytochrome oxidase I (COI) locus (Barth *et al*. 2006). COI is generally considered a ‘neutral’ marker with regard to traits under selection, such as infection genotype in our case. DNA extractions, PCR amplification and Sanger sequencing were performed as detailed either in (Barth *et al*. 2006) using primers COXL and COXH, or in (Struder-kypke 2010) using primers F199dT-B and R1143dT. An intraspecific COI phylogeny was constructed by aligning the sequences and constructing both Neighbor Joining and Maximum Likelihood phylogenies in the Geneious program (https://www.geneious.com/home/). Pairwise genetic distances across all strains and also within the A and B clades were calculated in Geneious and in Mega7 (https://www.megasoftware.net (Kumar *et al*. 2016).

### 2.5 Statistical analysis

To analyze variation in infection prevalence, generalized linear mixed effect models (GLMMs; binomial error structure, logit link) were carried using the SAS 9.4 software (SAS Institute Inc. 2013. *SAS/STAT 13*.*1 User’s Guide*. Cary, NC: SAS Institute Inc.). In a first analysis, strain identity and time point (early / late) were taken as fixed effects, and laboratory (LB / OK), experimental block and experimental replicate as random factors. In a second analysis, clade (A / B) and time point were taken as fixed effects, with strain, laboratory and experimental block as random factors. Preliminary analyses indicated no significant difference between day 5 and day 7 p.i. measurements in the LB lab (p > 0.3; correlation: r = 0.58, n = 18), and thus the data were combined as ‘early’ estimates in the analyses.

To quantify genetic variation in resistance, we calculated the broad-sense heritability, generally defined as the fraction of genetic variation over the total phenotypic variation in a trait (Falconer 1989). For this kind of data (clonal replicate cultures of a given strain), heritability is the intraclass correlation coefficient (ICC), i.e., the degree of similarity of resistance for replicates of the same strain. The ICC was calculated for early and late measurements of infection prevalence, across all strains combined and separately for clade A and B strains. To standardize the data, we first fitted a generalized linear model (GLM), with laboratory and experimental block as factors. On the residuals of this GLM, we performed one-way ANOVAs with strain as a random factor to obtain the variance components and to calculate the ICC (Zar 2013) and confidence intervals (Demetrashvili *et al*. 2016). Finally, we also performed Mantel tests to test for correlations between pairwise genetic, geographic and phenotypic distance matrices using the “vegan” package in R (Oksanen 2019). We further tested for a correlation between infection prevalence and geographical distance between host and parasite strains.

## 3 Results

### 3.1 *Paramecium caudatum* genotyping and geographic origin

Of the 30 COI sequences obtained from the *P. caudatum* strains, 26 represented genetically distinct genotypes (with three different genotypes represented by multiple strains). Phylogenetic analysis revealed that most strains fall into two main clades, defined by the A and B haplogroups (Figure 2, Table 1). The average pairwise distance at the COI locus among all strains genotyped was 0.039 (i.e., 3.9% distance between strains), which is consistent with genetic distances between known reproductively isolated species in other protists (Foissner 2009). The average pairwise distances within haplogroups A and B were about four times smaller than the overall average, reflecting a clear genetic separation between the two haplogroups. Within-group levels of COI diversity were similar for the A and B groups (0.007 and 0.009, respectively). The two strains representing the C and D haplogroups were both clear outgroups to the A and B clades.

**Table 1.** Average pairwise genetic distance and percent identity across COI sequences (626 nucleotides) across all 30 *Paramecium caudatum* strains and within the A and B haplogroups. Number of strains indicated in parentheses for each calculation. Aligned using Muscle and genetic distances calculated in Mega7, using the pairwise distance function. Identical sites determined in Muscle.

|  | average<br>pairwise<br>distance | % identical<br>sites across<br>sequence |
| --- | --- | --- |
| All strains (30) | 0.042 | 85.0% |
| Haplogroup A (20) | 0.009 | 95.0% |
| Haplogroup B (8) | 0.007 | 98.2% |

**Figure 2.**
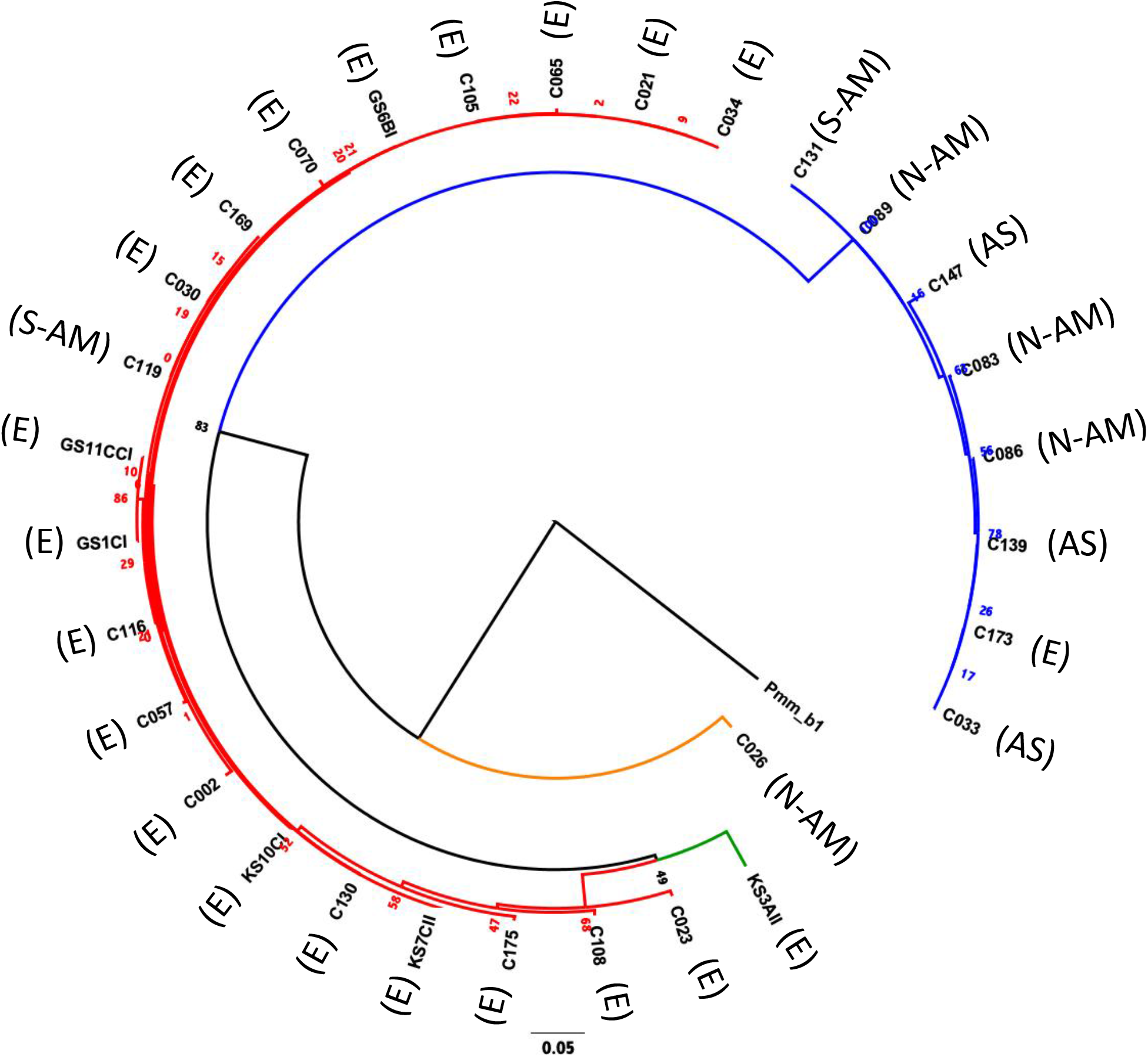
Maximum likelihood phylogeny (Tamura-Nei substitution model) constructed by comparing the COI sequence (626 nts) of the 30 *Paramecium caudatum* strains used in this study, using for an outgroup a *P. multimicronucleatum* isolate PmCOI_b1_01 (accession number AM072765.1) (Barth *et al*. 2006). Red branches = haplogroup A; blue branches = haplogroup B; orange branches = haplogroup C; green branches = haplogroup D. Geographic origins are as follows: E = European origin; AS = Asia; N-AM = North America; S-AM = South America. The number represent maximum likelihood estimates of branch lengths (mean number of substitutions per site).

There was a clear association between haplogroup identity and geographic origin. Notably, 19 out of 20 strains from haplogroup A had been collected in Europe, while B strains were of diverse (mostly non-European) origin (Figure 2, Table S1). This segregation between A and B haplogroups generated an overall positive correlation between pairwise genetic distances (COI sequence differences) and geographic distances (Mantel test: r = 0.45, p < 0.0001; Figure S2B). However, there was considerable variation around this pattern: strains with identical COI genotypes had very different geographic origins, and at a smaller scale, within Europe, the genetic-geographic correlation was weak (Mantel test: r = 0.15, p > 0.1).

### 3.2 Quantitative genetic variation in resistance

We observed substantial variation in resistance among the 30 *P. caudatum* strains tested (Figure 3, Figure S1), with results being highly repeatable between laboratories (correlation of strain means: r = 0.75, n = 16, p = 0.0008). Variation was essentially quantitative, with only one strain (C175) showing no sign of infection in any of the inoculated replicates. For the other strains, mean levels of infection ranged from 2.8% (strain C057) to 78% (strain C021). Thus, statistical analysis revealed a significant strain effect (Table 2A) and high values of broad-sense heritability of resistance (intraclass correlation coefficient > 0.5; Table 3).

**Table 2.**
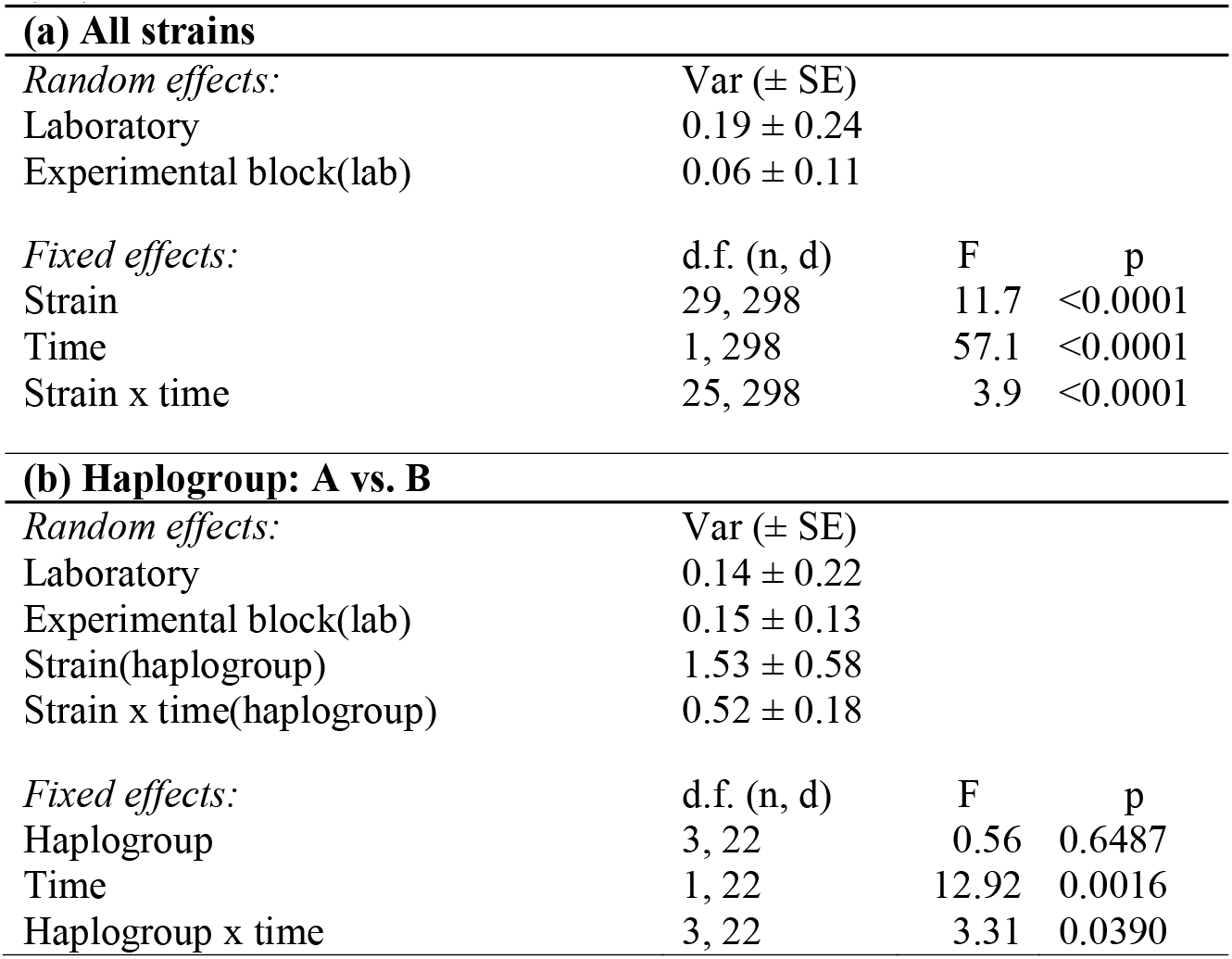
ANOVA results from GLMM models of resistance of *Paramecium caudatum* to *Holospora undulata*, as a function of strain identity, time (early vs. late measurements) and haplogroup (A vs. B). For fixed effect F-tests, numerator (n) and denominator (d) degrees of freedom are given. In all models, laboratory and experimental block were fitted as random factors, and their variance and standard error (SE) are given.

**Table 3.** Broad-sense heritability of resistance of *Paramecium caudatum* to *Holospora undulata*, for all strains combined and for strains from haplogroups A and B, respectively. Heritabilities represent intraclass correlation coefficients, based on measurements of infection prevalence one week (early) and 2-3 weeks (late) post inoculation. Confidence intervals (95%) shown in square brackets, the number of strains used for calculations in round brackets. Calculations based on residual infection prevalences, after correction for laboratory and block effects.

|  | Early | Late |
| --- | --- | --- |
| All strains | 0.80 [0.69; 0.87] (30) | 0.34 [0.21; 0.47] (26) |
| Haplogroup A | 0.80 [0.57; 0.84] (20) | 0.30 [0.15; 0.45] (17) |
| Haplogroup B | 0.75 [0.45; 0.90] (8) | 0.50 [0.21; 0.75] (7) |

**Figure 3.**
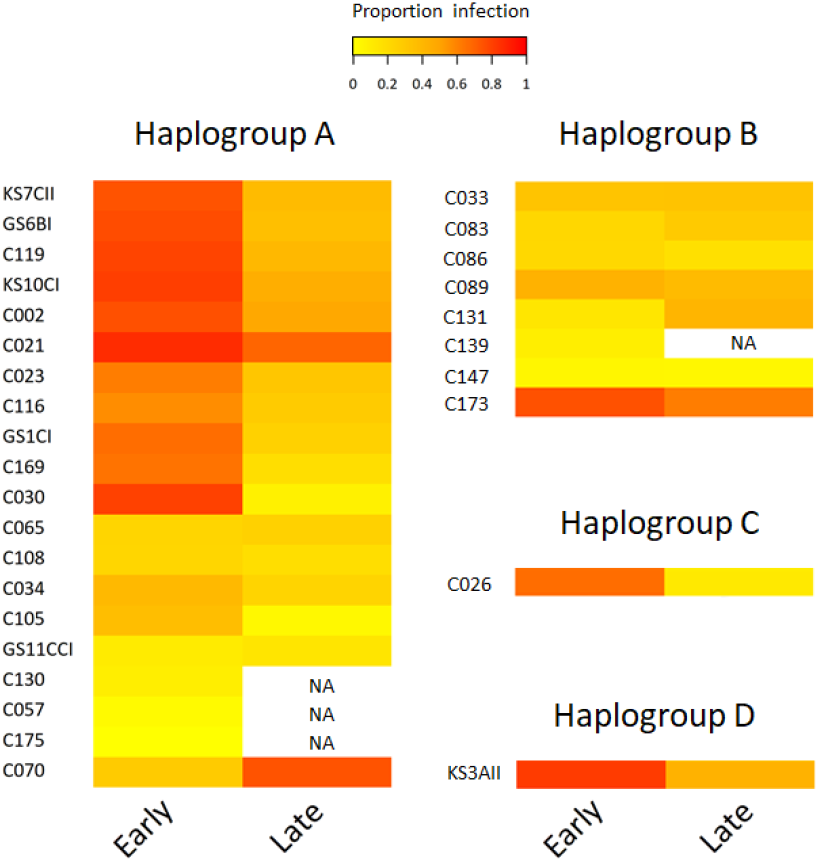
Heatmap of resistance of 30 *Paramecium caudatum* strains (from 4 COI haplogroups) against the parasite *Holospora undulata*. Measurements of infection prevalence (proportion of infected host cells) were taken 5-7 days (‘early’) and 2-3 weeks (‘late’) post inoculation. Each row represents a different strain. The color gradient, from yellow to red, indicates increasing levels of infection prevalence, and thus decreasing resistance.

#### 3.2.1 Variation by time point

We observed a significant general decrease in infection prevalence between the early and late time points (48 ± 2% vs 34 ± 2% SE). Although estimates of strain mean tended to be positively correlated between time points (r = 0.36, n = 26, p > 0.06), the significant time x strain interaction (Table 2A) indicates that differences between strains varied substantially according to time point (Figure 3). In at least two strains (C030, K3AII), there was a very strong decline in infection prevalence from initially high levels of infection to very low levels (<10%), with infection undetectable in several replicates.

Although prevalence declined in the majority of strains, it remained unchanged or even increased in some cases. In the case of C070, initially low levels rose to high levels of detectable infection, although we measured only one trial for this strain. For the majority of strains, however, we found increasing variation among replicates from the early to late timepoints (Figure S1), leading to a decline in heritability estimates by the late timepoint (Table 3). Thus, the amount of detectable genetic variation in resistance tended to lessen over the infection time course.

#### 3.2.2 Variation by haplogroup

Haplogroup A strains on the whole tended to have higher levels of infection (48.8 % ± 6.7 SE) than haplogroup B strains (27.8 % ± 8 SE, Figure 3), namely at the early time point (significant haplogroup x time point interaction, Table 2B). However, both haplogroups harbor significant and similar levels of genetic variation in this trait (Table 3). The two single strain representatives of haplogroups C and D were not included in the statistical analysis but were both highly susceptible to infection (Figure 2).

Aside from the overall differentiation between the A and B haplogroups, additional analyses provide little evidence for a link between COI genotype and resistance (Mantel test comparing genetic distances based on COI genotype and phenotypic differences in resistance: r = −0.05, p > 0.8; Figure S2C). Indeed, strains with identical COI genotypes had widely varying resistance phenotypes (C105 & GS6BI; C030 & C119; C033 & C086 & C139 & C173; see Figure 3).

#### 3.2.3 Variation by geographic area

Although haplogroup identity coincided in part with large-scale geographic differentiation (Europe vs rest of world), we found no significant overall relationship between pairwise geographic distances and differences in resistance (Mantel test: r = −0.02, p > 0.6; Figure S2A). However, at the more regional level, within Europe, geographically closer strains tend to have more similar levels of resistance (Mantel test: r = 0.16, p = 0.0487; Figure 4A). In addition, the probability of infection of European strains was negatively correlated with their geographic distance from the site of origin of the parasite (r = −0.625, p = 0.002): strains that are geographically closer to the parasite location were more likely to become infected than those from further away (Figure 4B).

**Figure 4.**
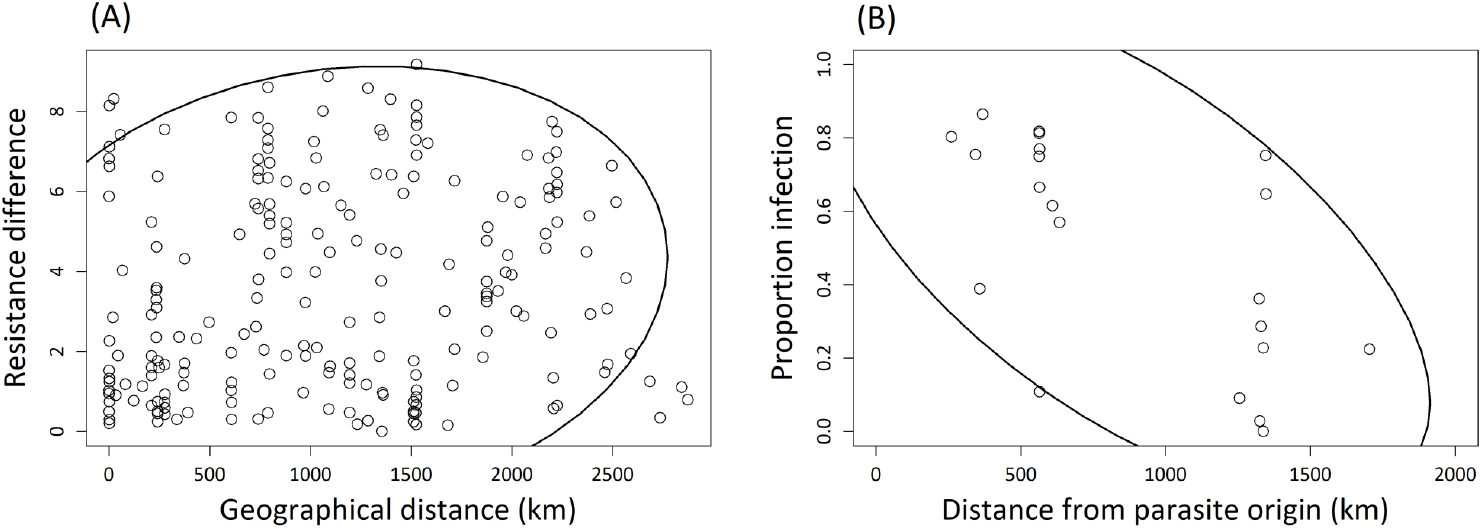
**A**. Correlation between pairwise geographic Euclidean distances and pairwise resistance differences, for 21 European *Paramecium caudatum* strains inoculated with the parasite *Holospora undulata*. Resistance differences based on the mean residual infection prevalences for each strain, after statistically correcting for laboratory and experimental block effects. Absolute values of pairwise differences were used for the correlation analysis. Each point refers to a different pair of strains. **B**. Correlation between infection prevalence and the geographic distance between the origin of a given *P. caudatum* strain and the origin of parasite tester isolate. Infection prevalence was averaged over multiple replicates per strain. Each point represents one strain. The 90% density ellipsoids represent a graphical indicator of the correlation between the two variables in each panel.

## 4 Discussion

### 4.1 High levels of heritability: Resistance is a quantitative trait with a genetic basis

Resistance is a key determinant in interactions between hosts and their parasites, and studying the amount and distribution of the genetic variation in this trait can provide insights in the possible (co)evolutionary processes in natural populations (Thompson 2014). While previous work described qualitative patterns of resistance variation in the *Paramecium-Holospora* system, we went one step further by investigating the quantitative genetic variation in resistance to *H. undulata* across a collection of *P. caudatum* strains, using controlled inoculation experiments. We found substantial levels of heritability for resistance, in a pattern resembling a continuum from total resistance (0% infection prevalence) to near-complete susceptibility (80%) of strains.

From a quantitative genetics perspective, this result indicates that there is a genetic basis of resistance upon which selection can act in natural populations. Despite a somewhat artificial experimental setup, using inocula with large amounts of transmission stages, all transmission occurred naturally in our tests, and results were repeatable between laboratories and consistent over independent experimental replicate runs. Moreover, genetic differences in resistance persisted even over multiple asexual generations in our experimental cultures, where demographic and epidemiological processes acted freely. In the longer run (several months), we can detect increased levels of *de novo* resistance in such cultures, e.g., (Duncan *et al*. 2011c), demonstrating that this trait indeed responds to parasite-mediated selection.

If *Holospora* truly selects for increased resistance, we need to explain why so much natural genetic variation is still present in this trait. One possibility is variation in the strength of parasite-mediated selection. Some of our strains may originate from unexposed populations and have no (co)evolutionary history with the parasite, which would explain the finding of highly susceptible genotypes in the collection. This is plausible, given the low incidence reported for *Holospora* infections (Fokin 2009). Moreover, even if *Holospora* were more frequent, high-resistance variants may not necessarily spread to fixation if resistance is associated with a cost. Indeed, trade-offs between resistance and other fitness-relevant traits are common in many host-parasite systems (Duncan *et al*. 2011a). For the present collection of strains, higher equilibrium density (when uninfected) tends to be associated with higher susceptibility (r = 0.22, n = 29, p > 0.2), indicative of a weak cost of resistance, already previously documented for our system (Lohse *et al*. 2006; Duncan *et al*. 2011a; Duncan *et al*. 2011c).

### 4.2 Specificity: evidence for haplogroup differences and local adaptation?

Another potential mechanism contributing to the maintenance of genetic diversity is genotype-specific adaptation, or more generally, host genotype x parasite genotype interactions (Lambrechts *et al*. 2005; Cayetano and Vorburger 2013). Previous work indicated large-scale differences in qualitative resistance between entire syngens, or mating groups, within *Paramecium* species (Fujishima and Fujita 1985; Rautian 1993; Skoblo 1996; Potekhin *et al*. 2018). This might reflect signatures of lineage-specific adaptation, such that certain groups of *Holospora* strains can only infect certain *Paramecium* syngens (Rautian 1993). In our study, we did not find such a qualitative difference in resistance among the four highly divergent *P. caudatum* haplogroups, i.e., all haplogroups tested could be infected. However, the A and B haplogroups showed a general difference in mean resistance levels, at least at the early time point (Figure 3).

The A and B haplogroups can be broadly divided into European (A) and non-European (B) strains. We speculate that the *Holospora* strain that we used as a tester, which is of European origin, might have an evolutionary history with haplogroup-A strains and is therefore, on average, better adapted to infect strains with this same European genetic background. This is consistent with observations at a finer resolution. First, there was a weak signal of geographic distance among European strains, such that more closely located strains were more similar in their resistance phenotype (Figure 4A). This indicates that the spatial distribution of resistance follows geographic patterns of isolation by distance. Second, and more importantly, we found a negative correlation between resistance and geographic distance between parasite and host strains (Figure 4B). This distance effect likely reflects a signature of parasite local adaptation (Ebert 1994; Kaltz 1998; Kaltz *et al*. 1999), with infection success of our tester strain being highest on host genotypes that are geographically close to its origin (South of Germany). These interpretations are based on tests with a single parasite isolate. A thorough assessment of these patterns requires reciprocal cross-infection assays, testing parasite isolates from different haplogroups for lineage specificity, or testing sympatric and allopatric combinations of host and parasite genotypes for local adaptation.

Infection experiments can profit immensely from accompanying molecular analysis of the geographic host (or parasite) population structure (Parratt 2016; Andras *et al*. 2018). Our work shows that a single conserved neutral marker such as COI can be used for large-scale differentiation of lineages but is of limited utility in distinguishing closely related strains from one another. We have several instances among our strains where the COI genotypes of different strains are identical, but their infection phenotypes are divergent (Figure 3). Obviously, other loci in the genome control this trait and distinguish these different strains from one another. Future whole-genome sequencing efforts and association studies promise to untangle these genotype-phenotype interactions more definitively.

### 4.3 Mechanisms of resistance: quantitative variation caused by multiple infection barriers

Our results clearly show that resistance variation is quantitative, or perhaps more precisely, probabilistic: upon inoculation, only a certain fraction of the inoculated hosts become infected. This might be simply due to random effects related to the number of cells ingesting IFs, but could also be more complex, for example if multiple host factors (and possibly interactions with multiple parasite factors) or a multi-step process (perhaps involving multiple signaling steps) determine infection success (Agrawal and Lively 2003; Hall *et al*. 2019). It appears that nearly all *P. caudatum* strains examined here are potentially ‘colonizable’ in that the defenses could be breached, allowing the parasite to enter the cell and invade the nucleus. Whether colonization occurs may depend on the efficiency of the host signaling and surveillance; this would then determine the probability of infection at the individual level and translate into a given population-level infection prevalence.

Resistance mechanisms in the host can be deployed at different points during the first steps of the infection process, including selective uptake of IFs by the host, inability of the IF to break out of the food vacuole or to enter the nucleus, or lysis in the nucleus at early stages of bacterial development (Rautian 1993; Fokin 1997; Skovorodkin 2001; Fokin 2005; Fels *et al*. 2008; Goertz 2010). These effects should have occurred before our ‘early’ measurement timepoint of infection prevalence. Therefore, variation at loci controlling these mechanisms could provide the quantitative genetic variation that we observed in these experiments at the early timepoint. The relative contribution of different resistance mechanisms could be investigated through repeated sampling at very early time points post inoculation, from which we could track uptake rates of IFs, the frequency and number of IFs entering the nucleus, etc (Fels and Kaltz 2006; Fels *et al*. 2008).

Previous work reported strain-specific drops in infection levels or total loss of infection over longer time spans, evoking additional action of ‘late resistance’ (Ossipov *et al*. 1993; Skoblo 1996; Fokin 1997; Fokin *et al*. 2003). Our study confirms a general trend of decline in infection prevalence 2-3 weeks post inoculation, at times to very low levels (<10%) or even below detection thresholds (see Figure S1). However, these observations can also be explained by contributions from other factors, such as the fidelity of vertical transmission, the parasite’s virulence or its investment into horizontal transmission (Nidelet *et al*. 2009; Magalon *et al*. 2010), jointly affecting the demography and epidemiology in the culture. Variation in these parameters can be estimated from time-series data and the use of epidemiological models (Lunn *et al*. 2013; Nørgaard 2020).

### 4.4 Conclusions and future directions

Our study provides a quantitative assessment of the amount and distribution of natural genetic variation in resistance in the *Paramecium* - *Holospora* system and combines available information on the phylogeny and biogeography with the observed phenotypic variation. We demonstrate ample heritable variation in resistance on which selection can act and find tentative evidence of symbiont adaptation producing signatures in geographic and/or clade-specific patterns of resistance, typical of many host-parasite systems (Thompson 2005).

As mentioned above, more extensive cross-inoculation tests, sampling both host and symbiont across multiple geographic locations and haplogroups, will be required to obtain a better picture of the evolutionary forces shaping the observed variation. Such approaches can be complemented by cross mating *Paramecium* strains in standard diallel designs (Falconer 1989) and measuring resistance in the offspring. This would eliminate potential strain-specific epigenetic or somatic variation arisen in the macronucleus (Beales 2008) not accounted for in the present experiment. Furthermore, combining phenotypic assays with genomic studies, both whole genome sequencing and RNA sequencing over a time course, will help to identify particular gene variants involved in infection, and in so doing, point the way to the precise molecular mechanisms involved, as well as the evolutionary patterns across geography and phylogeny.

## 6 Supplementary figure legends

**Figure S1.**
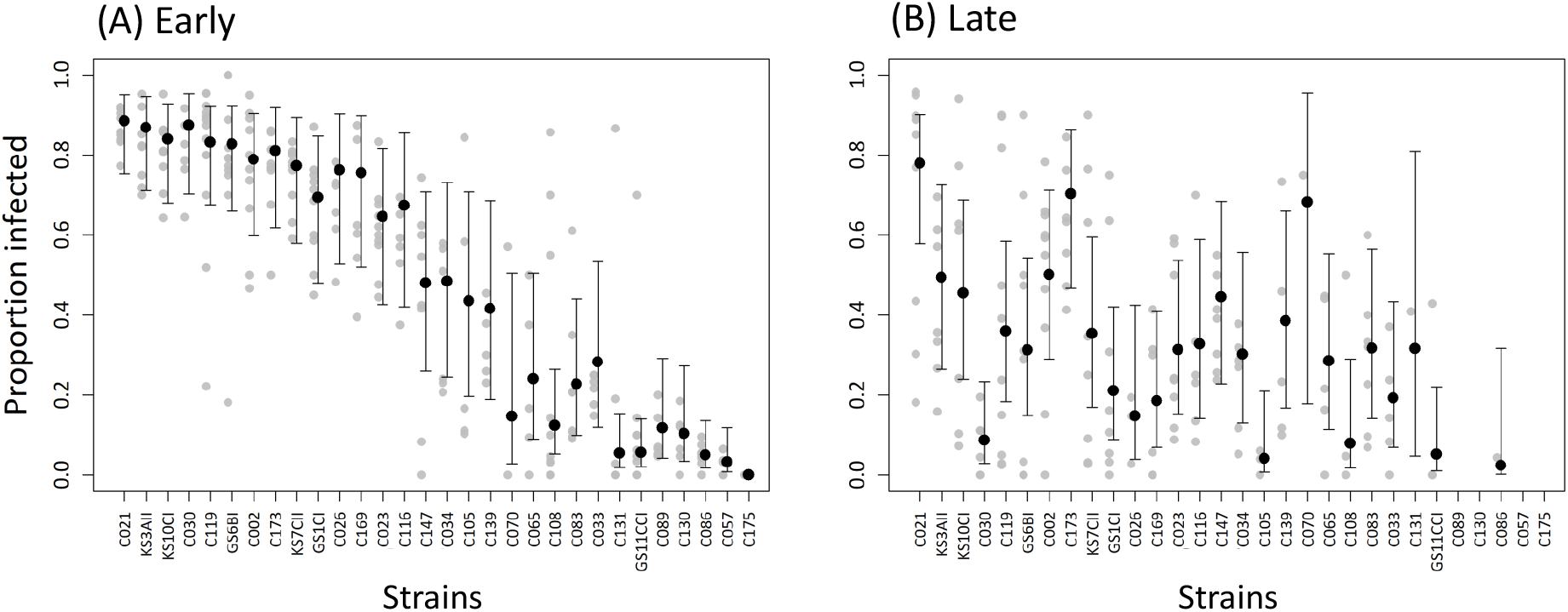
Infection prevalence in 30 *Paramecium caudatum* strains, as measured at **(A)** early time points (5 / 7 days) and **(B)** late time points (14 / 20 days) after infection with *Holospora undulata*. Light grey points are measured data, while the bigger black points are the model predictions of the statistical analysis with 95% confidence interval.

**Figure S2.**
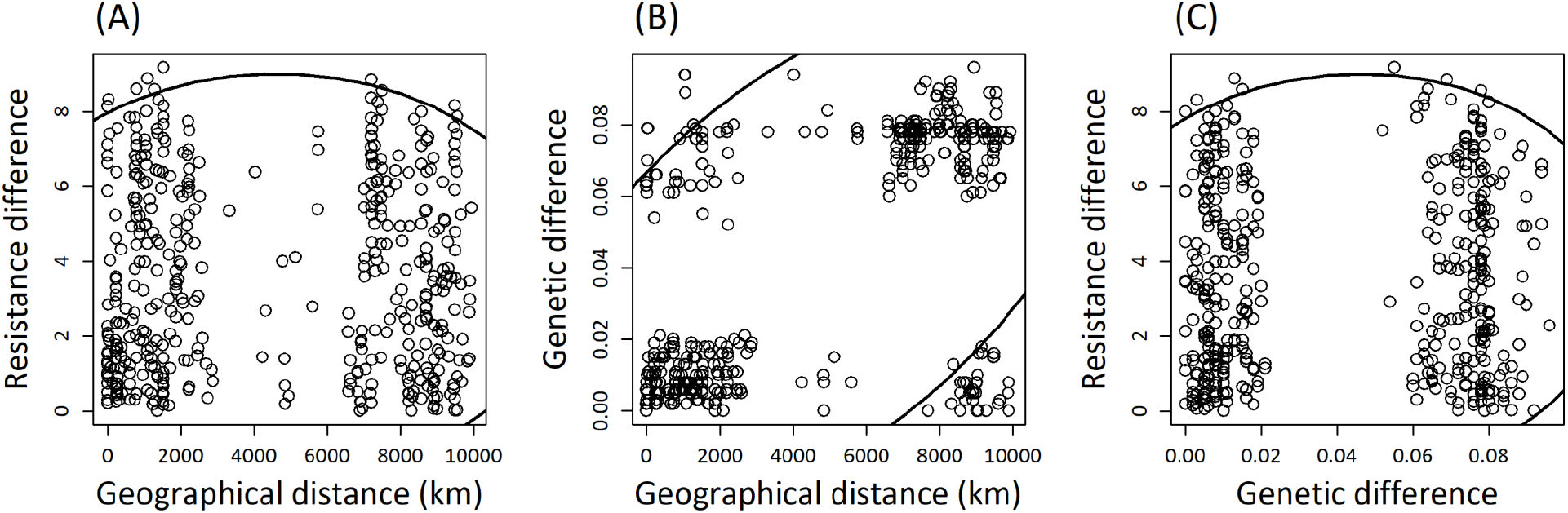
Correlations between pairwise genetic distances (based on neutral COI marker), geographic Euclidean distances and pairwise resistance differences, for 30 *Paramecium caudatum* strains confronted with the parasite *Holospora undulata*. Resistance differences based on the mean residual infection prevalences for each strain, after statistically correcting for laboratory and experimental block effects. Absolute values of pairwise differences were used for the correlation analysis. Each point refers to a different pair of strains. The 90% density ellipsoids represent a graphical indicator of the correlation between the two variables in each panel.

## 8 Supplementary tables

**Table S1**. Information on the 30 *Paramecium caudatum* strains used in this study, including geographic origin and year of sampling, cytochrome oxidase 1 (COI) genotype and haplogroup, and the number of replicates tested in the resistance assays in the Bright (LB) and Kaltz (OK) labs.

## 10 Author Contributions

LJB and OK conceived and designed the experiments; JW, GZ, SK, NZ, LN, and WC collected the data; JW, GZ, SK, LJB, and OK analyzed and interpreted the data; LJB, JW, GZ, and OK drafted the article; LJB, GZ, LN, and OK performed critical revision of the article; all authors gave final approval of the version submitted for publication.

## 12 Acknowledgments

We thank Sebastien Tarcz for providing several unpublished COI sequences of strains, including Hap97, Hap98, Hap99, and Hap100. From the Bright lab, Nicole Lee and Avery Weiner contributed to early infection tests and cell culture maintenance. Claire Gougat-Barbera contributed to culture maintenance in the Kaltz lab.

