## Supplementary material for "Heritable variation in resistance to the endonuclear parasite *Holospora undulata* across clades of *Paramecium caudatum*": Table S1.

Table S1. Information on the 31 *Paramecium caudatum* strains used in this study, including geographic origin and year of sampling, cytochrome oxidase 1 (COI) genotype and haplogroup, and the number of replicates tested in the resistance assays in the Bright (LB) and Kaltz (OK) labs. (*) indicates approximate location.

| Strain | Location code | Origin | GPS coord. | Year | COI genotype | COI haplogroup | Replicates  (LB / OK) |
| --- | --- | --- | --- | --- | --- | --- | --- |
| C002 | Hainberger See | Germany, Saxony | 51.035688, 12.285028 | 2005 | PcCOI_a12 | A | 3 / 6 |
| C021 | Erzgebirge E1 | Germany, Saxony | 50.647032, 13.256963 | 2009 | PcCOI_a14 | A | 3 / 6 |
| C034 | Frohburg 1 | Germany, Saxony | 51.050148, 12.557741 | 2006 | PcCOI_a27 | A | - / 6 |
| C023 | Plön K 2 | Germany, Schleswig-Holstein | 54.15021, 10.440307 | 2005 | PcCOI_a19 | A | 3 / 6 |
| GS11CCI | Globsowsee | Germany, Brandenburg | 53.128590, 13.118740 | 2017 | Hap97 (**) | A | 3 / 6 |
| GS1CI | Globsowsee | Germany, Brandenburg | 53.128590, 13.118740 | 2017 | Hap97 (**) | A | 2 / 6 |
| GS6BI | Globsowsee | Germany, Brandenburg | 53.128590, 13.118740 | 2017 | PcCOI_a01 | A | 3 / 6 |
| KS10CI | Kochsee | Germany, Brandenburg | 53.131996, 13.107747 | 2017 | Hap98 (**) | A | 3 / 6 |
| KS7CII | Kochsee | Germany, Brandenburg | 53.131996, 13.107747 | 2017 | Hap100 (**) | A | 2 / 6 |
| C030 | Österreich_Lahnalp | Austria, Tyrol | 47.695948, 12.218064 (*) | 2005 | PcCOI_a03 | A | - / 6 |
| C057 | SWE 12.1 | Sweden, Dalarnas län | 60.128605, 15.169305 | 2005 | PcCOI_a25 | A | - / 6 |
| C065 | SWE 17.1 | Sweden, Dalarnas län | 60.105747, 15.966911 | 2005 | PcCOI_a26 | A | - / 5 |
| C130 | SWE 9.2 | Sweden, Värmlands län | 59.790063, 13.067001 | 2005 | PcCOI_a24 | A | - / 6 |
| C070 | Norw 8 Gol | Norway, Viken | 60.700700, 8.978653 (*) | 2005 | PcCOI_a16 | A | 1 / - |
| C105 | Sp 10C | Spain, Valencian Community | 39.548852, -1.502887 | 2005 | PcCOI_a01 | A | - / 5 |
| C108 | Portugal A1 | Portugal, Alentejo | 38.776742, -7.171436 | 2005 | PcCOI_a28 | A | 3 / 6 |
| C116 | Frankreich 10-2.1 | France, région PACA | 43.430297, 6.126616 | 2006 | PcCOI_a30 | A | - / 6 |
| C169 | Greece 2.1 | Greece, Central Macedonia | 41.007458, 22.2761 | 2006 | PcCOI_a31 | A | - / 6 |
| C175 | Greece 11 | Greece, Western Macedonia | 40.709844, 21.744039 | 2006 | PcCOI_a32 | A | - / 6 |
| C119 | Peru | Peru, ? | -12.046184, -77.040842 (*) | 2006 | PcCOI_a03 | A | 4 / 6 |
| C033 | Peking 1_C3 | China, Beijing | 39.128165, 117.185083 (*) | 2006 | PcCOI_b01 | B | - / 6 |
| C083 | USBL-5I1 | USA, Indiana | 39.069861, -86.414361 | 2011 | PcCOI_b12 | B | 1 / 6 |
| C086 | USBL-11II6 | USA, Indiana | 39.179472, -86.534333 | 2011 | PcCOI_b01 | B | 1 / 6 |
| C089 | YE1I | USA, Indiana | 39.196563, -86.511732 (*) | 2011 | PcCOI_b013 | B | - / 6 |
| C131 | Equador 17 III | Ecuador, Loja | -4.018924, -79.202980 (*) | 2010 | PcCOI_b11 | B | 1 / 6 |
| C139 | My43c3d | Japan, ? | 38.480973, 141.372414 (*) | 2010 | PcCOI_b01 | B | - / 5 |
| C147 | KNZ5414 | Japan, Ishikawa Prefecture | 36.519469, 136.709415 | ? | PcCOI_b07 | B | 1 / 6 |
| C173 | Greece 10.1 | Greece, Central Macedonia | 40.805947, 21.983306 | 2006 | PcCOI_b01 | B | 1 / 6 |
| C026 | Fokin 1 UBR 4-2(Baton Rouge) | USA, Louisiana | 30.46788, -91.129604 (*) | ? | PcCOI_c02 | C | - / 6 |
| KS3AII | Kochsee | Germany, Brandenburg | 53.131996, 13.107747 | 2017 | Hap99 (**) | D | 1 / 6 |
